# Characterisation of ligand gating, ion conduction and the ion selectivity mechanism in the endo-lysosomal ion channel hTPC2

**DOI:** 10.1101/2023.11.01.565142

**Authors:** Alp Tegin Şahin, Ulrich Zachariae

## Abstract

Two pore channels (TPCs) are two-fold symmetric endo-lysosomal cation channels forming important drug targets especially for antiviral drugs. They are activated by calcium, ligand binding, and membrane voltage, and to date, are the only ion channels shown to alter their ion selectivity depending on the type of bound ligand. However, despite their importance in the field, ligand activation of TPCs and the molecular mechanisms underlying their ion selectivity are still poorly understood. Here, we set out to elucidate the mechanistic basis for the ion selectivity of human TPC2 (hTPC2) and the molecular mechanism of ligand-induced channel activation by the lipid PI_(3,5)_P_2_. We performed all-atom in silico electrophysiology simulations to study Na^+^ and Ca^2+^ permeation across hTPC2 in real-time and to investigate the conformational changes induced by the presence or absence of bound PI_(3,5)_P_2_. Our findings reveal that hTPC2 adopts distinct structures depending on the presence of PI_(3,5)_P_2_ and elucidate the conformational transition pathways between these structures. Additionally, we examined the permeation mechanism, solvation states, and binding sites of ions during ion permeation through the pore. Our simulations reproduce the experimental observation that hTPC2 is more selective for Na^+^ over Ca^2+^ ions in the presence of PI_(3,5)_P_2_ and explain the mechanism of this ion selectivity. They highlight the importance of specific ion binding sites at the luminal channel entrance, the selectivity filter, and the central channel cavity for ion conduction, enabling a distant knock-on mechanism for efficient permeation of Na^+^ ions.

## Introduction

Ion channels are essential components of all cells. They form membrane-integral proteins that mediate efficient and often selective ion exchange across lipid membranes, which are otherwise impermeable to ions. Their function is key to maintaining ionic homeostasis in the cell and its compartments, as well as constituting one of several ways for a cell to communicate with its surroundings, allowing cells to use electric currents to transmit information and share it with neighbouring cells^[1]^. Inside eukaryotic cells, organelles employ similar principles to regulate intracellular ion exchange and ion-based communication, facilitated by the function of organellar and vesicular ion channels.

Two-Pore Channels (TPCs) form an important channel family found primarily in endo-lysosomal membranes^[2]^. Endo-lysosomes are membrane-enclosed organelles, which play a key role in cell survival and cell death. They are also crucial for viral entry into and release from eukaryotic host cells^[3]^. For this reason, TPCs represent prime targets for antiviral agents against viruses such as Ebola and SARS-coronaviruses. In the case of Ebola, TPCs have been validated as a major target for the development of antiviral drugs and drug design efforts are presently under way^[4,5]^

TPCs, comprising the subfamilies TPC1-3^[2,6]^, are ancient members of the cation-selective, voltage-gated ion channel superfamily, sharing sequence similarities with other types of eukaryotic Na^+^- and Ca^2+^-selective channels such as transient receptor potential (TRP) channels. They are believed to represent an evolutionary transition stage from the four-domain monomeric to the homo-tetrameric, more complex voltage-gated ion channels^[7]^. In accordance with their organellar location, encoding genes of TPCs can be found in eukaryotes but not in prokaryotes^[8]^. TPC1 is more abundant in endo-lysosomal organelles while TPC2 is preferentially localized to late endosomes and lysosomes^[9]^. TPC3 is not expressed in humans but mammalian and vertebrate orthologs can be found both in endo-lysosomal and plasma membranes^[6,9]^.

In the endo-lysosomal membrane, TPCs form homodimers. Despite their variety and the existence of different isoforms, the overall structure of TPCs is highly conserved^[10]^. Each subunit is comprised of two homologous Shaker-like six-transmembrane (TM) repeat domains (D1 and D2; Fig. 1)^[11]^. A single chain of TPCs thus contains two voltage-sensing domains (VSDs), formed by S1-S4 and S7-S10, as well as two pore-forming domains composed of helices S5-S6, S11-S12 (Fig. 1). The existence of two pore-forming domains per subunit has given rise to the nomenclature of this channel family^[11,12]^. In addition to their TM domains, TPCs possess EF-hand motifs on the cytosolic side, which are generally known as Ca^2+^ recruitment sites^[13]^. As opposed to plant TPC1, the mammalian TPCs’ EF-hands however lack the essential negatively charged residues, and as a result, Ca^2+^ is unlikely to bind the mammalian-type EF-hand motifs in TPCs^[14]^. Finally, the subunits comprise a C-terminal domain (CTD) and an N-terminal domain (NTD), which have been reported to play a functionally important role in channel activation. Deletion of these domains prevents the activity of the channel, but their mechanistic importance and role in the channels’ ion selectivity have yet to be elucidated^[11]^.

**Figure 1:**
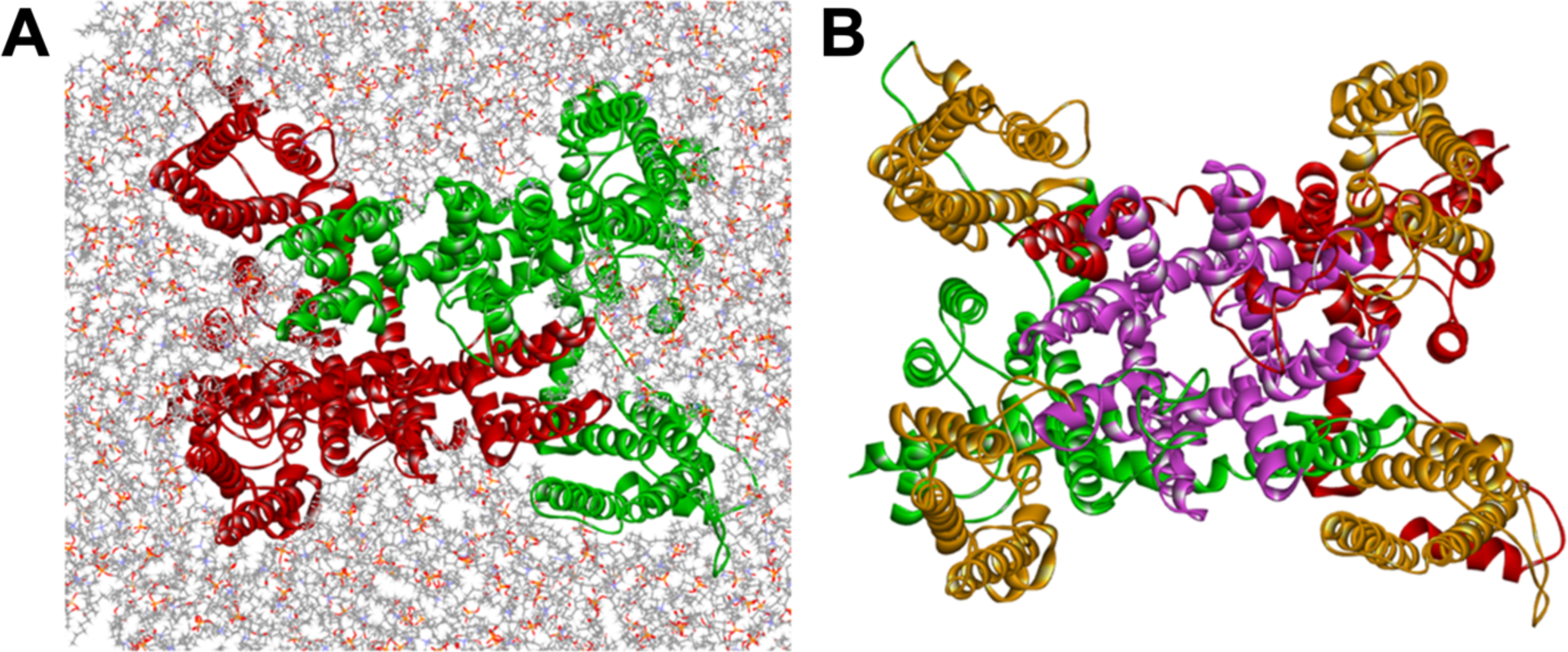
(A) Top view of the dimeric structure of human TPC2 (hTPC2) in a POPC membrane; the two subunits are shown in red and green. (B) The pore-forming domain (purple) and voltage-sensing domains (orange) are highlighted within the two subunits of hTPC2.

Channels of the TPC family can be gated by both voltage stimuli and a range of different ligands^[15]^. In particular, the channels respond to the binding of NAADP (nicotinic acid adenine dinucleotide phosphate) and the lipid PI_(3,5)_P_2_ (phosphatidylinositol-3,5-bisphosphate; PIP2)^[12,16]^, alongside electrochemical or voltage gradients. They are permeable to the cations Na^+^, Ca^2+^ and K^+^, each with different permeability levels (Fig. 2)^[17,18]^. TPC2 is known to be largely voltage-insensitive and activated mainly by ligand binding. In the PIP_2_-bound state, human TPC2 (hTPC2) is highly selective for Na^+^ compared to Ca^2+^, with a permeability ratio of P_Na_:P_Ca_ ≈ 10:1 in whole lysosomal recordings and P_Na_:P_Ca_ ≈ 17:1 in plasma membrane recordings^[16,19]^. In contrast, the activation of TPC1 is highly dependent on both voltage and ligands^[12,15,20,21]^.

**Figure 2:**
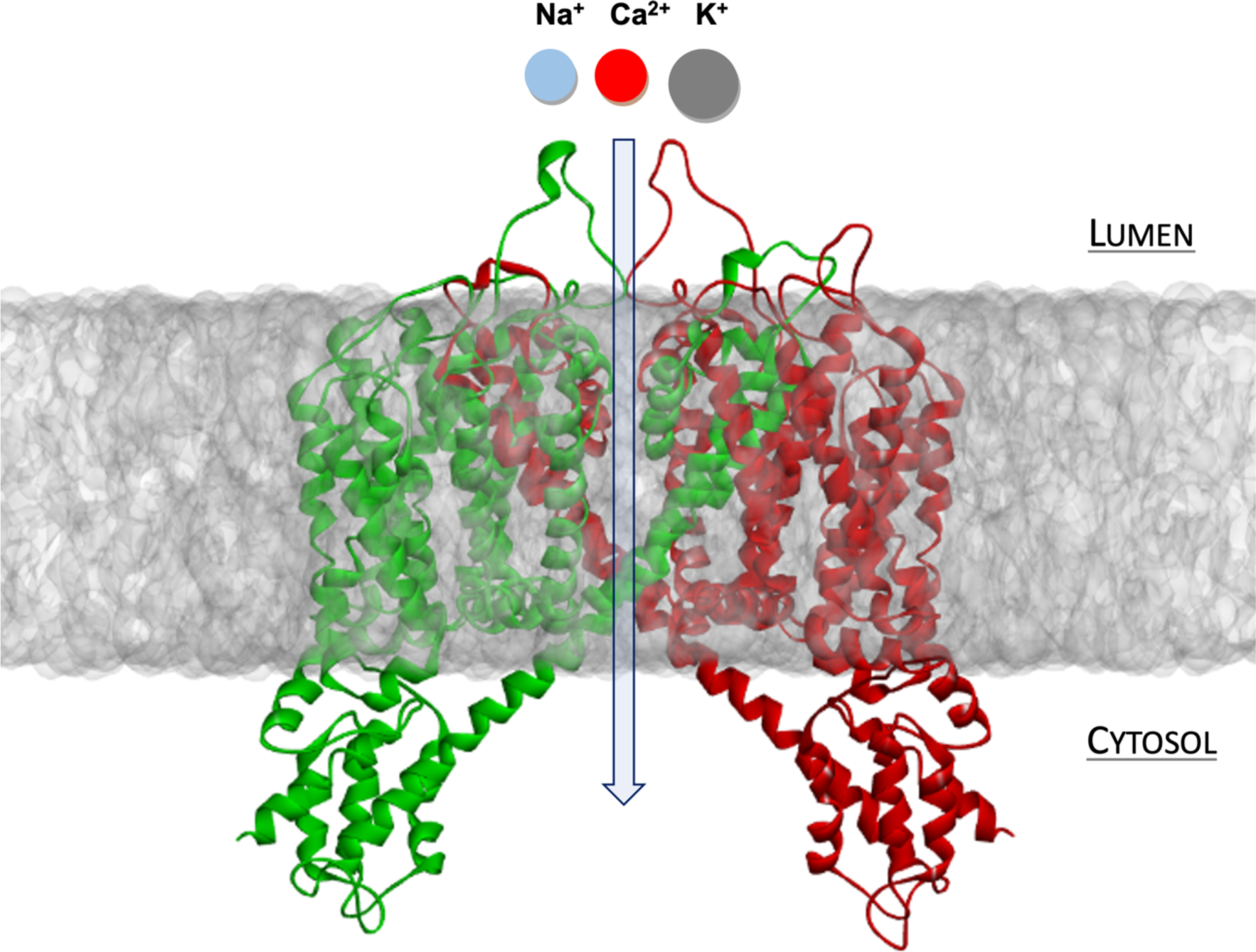
Transmembrane permeation pathway of cations through hTPC from the endo-lysosomal lumen to the cytosol (blue arrow) and translocated ion types. hTPC subunits are shown in red and green.

PIP2 trafficking regulates a wide range of cellular processes and is especially associated with endo-lysosomal functions. In the case of hTPC2, the binding of PIP2 alone is sufficient to activate the channels, with a variety of different studies showing that PIP2 binding triggers Na^+^ flux through TPCs^[22]^. Notably, recent studies have demonstrated that in the presence of NAADP, TPCs can change their ion selectivity from Na^+^ to Ca^2+^. This means that to date, TPCs are the only ion channels reported to change their ion selectivity in response to ligand binding^[12,15,22–25]^. NAADP is a potent and well-known Ca^2+^-mobilizing second messenger, which initiates Ca^2+^ release from endo-lysosomal organelles^[19]^. Recent research on the NAADP regulation of TPCs has shown that NAADP likely does not directly bind to the TPCs but via binding proteins such as Lsm12^[26][18,27]^.

So far, the selectivity and ligand-gating mechanisms of hTPC2 are insufficiently understood. The selectivity filter (SF) sequences of hTPC2 are _271_TAN_273_ (filter I) and _652_VNN_654_ (filter II)^[28]^. The hTPC2 cryo-EM structure shows that these two groups of SF residues are located at the entrance of the pore region^[16]^. These residues are conserved between the Na^+^-selective human and mouse TPC1-2 but not in the non-selective plant TPC1^[17]^. Recently, Guo et al. reported that mutations of V652 and N653 of filter II substantially affected the selectivity of hTPC2 for Na^+^, whereas mutation of A272 only had a subtle effect, indicating that the residues on filter II may be the main determinants of TPC selectivity^[28,29]^. Furthermore, molecular dynamics (MD) simulations of TPC1 and TPC2 have demonstrated the importance of hydrophobic gating and the difference in overall protein flexibility between the apo and the holo (lipid-bound) channel states^[30]^. Additionally, they highlighted the role of the central channel cavity for Na^+^ interactions^[30]^.

Here, we set out to elucidate the mechanistic basis for ion selectivity of hTPC2 and the molecular mechanism of ligand-induced channel activation by PIP2. We used *in silico* electrophysiology simulations to study Na^+^ and Ca^2+^ permeation across hTPC2 in real time and investigated the conformational changes related to pore gating induced by the presence or absence of PIP_2_. Our findings show that the PIP_2_ interaction with the VSDs is allosterically linked to conformational changes in the SF as well as the cytoplasmic bundle crossing (hydrophobic gate). According to our simulations, ion selectivity arises from a multi-layered mechanism comprising the luminal channel entrance, the SF, and the central channel cavity, enabling distant knock-on of ions.

## Materials and Methods

Our study was based on the recently published cryo-EM structure of the PIP_2_-bound state of hTPC2, described as the channel open state (PDB ID: 6NQ0)^[16]^. hTPC2 was embedded into two different membrane compositions, mammalian lysosomal membranes (LysM) (Table S1) and 1-palmitoyl-2-oleoyl-sn-glycerol-3-phosphocholine (POPC) membranes. Lysosomal and POPC bilayer systems were generated using the CHARMM-GUI server^[31]^. The initial box had a dimension of 150 × 150 Å in x and y-direction. The protein was aligned in the bilayer by using the PPM server^[31,32]^.

Unless otherwise stated, we carried out five repeat simulations for each different setup and simulation condition investigated. In each condition, the system was equilibrated initially for 2 ns, then a 20 ns pre-production equilibration without membrane voltage was performed, and following this, different lengths of production runs (between 250 ns and 500 ns duration each) with or without membrane voltage were carried out. Membrane voltages were generated using an applied external electric field yielding voltages of +700 mV, -700 mV, -350 mV, or -200 mV, respectively^[33]^. To examine the ion selectivity of hTPC2 in aqueous solutions, we used either 0.6 M NaCl or 0.6 M CaCl_2_ in mono-cationic solutions, or a mixture of 0.3 M NaCl and 0.3 M CaCl_2_ in di-cationic solutions, together with the TIP3P model for water molecules^[34]^. Ions and water were added with GROMACS 2022, which was also used for all simulations^[35]^. The CHARMM36m force field was used for the protein, ions other than Ca^2+^, and the lipids^[36]^. For the Ca^2+^ ions, the multi-site model developed by Zhang et al. was used, which has recently been shown by us and other groups to reliably model Ca^2+^ currents in a range of ion channels^[37–41]^

Hydrogen mass re-partitioning (HMR) of the system was used in the simulations with Na^+^ ions such that 4-fs time steps could be employed^[42]^. However, 4-fs time steps are incompatible with the multi-site Ca^2+^ model, and consequently, a standard 2-fs time step was used in all simulations with Ca^2+^, retaining HMR for consistency. For the investigation of the pore structure, architecture and radius profile, the CHAP program was used^[43]^. The LINCS algorithm was utilized to constrain bond lengths involving hydrogen atoms^[44]^. The systems were simulated in the NpT ensemble, where the temperature was maintained at T = 310 K using the Nosé-Hoover thermostat^[45]^, and the pressure was maintained semi-isotropically at 1 bar using the Parrinello-Rahman barostat^[46]^

## Results and Discussion

### Ligand-dependent opening of the channel at both the selectivity filter and the hydrophobic gate

We first performed simulations of hTPC2 in a POPC bilayer with and without bound PIP2. Both sets of simulations were initiated from the PIP2-bound cryo-EM structure of hTPC2 (PDB ID: 6NQ0), referred to as open-state structure in the PDB, with PIP2 bound to its binding pocket or removed, respectively. In the PIP2-bound simulations, we observed a substantial further dilation at the luminal SF region of hTPC2, increasing the dynamic pore diameter of the cryo-EM structure from ∼4-6 Å to ∼12-14 Å (Fig. 4). The key factor behind this widening was a rearrangement of both, the twin conserved asparagine residues N653 and N654 from filter II, and the twin conserved asparagine residues N273 and N274 from filter I on the neighbouring TPC2 subunit (Fig. 4A). Whereas the side chains of N653 and N654 point into the pore entrance in the cryo-EM structure, they reorientated tangentially to the pore in the simulations. Especially the side chains of N653 and N274 formed tight inter-subunit H-bonding interactions, dilating the luminal SF entrance. This super-open state was sustained in all simulations with bound PIP2 that showed the asparagine rearrangement at the SF entrance (6 out of 10 simulations in total). In the simulations without bound PIP2, by contrast, no such transition was observed at the SF.

**Figure 3:**
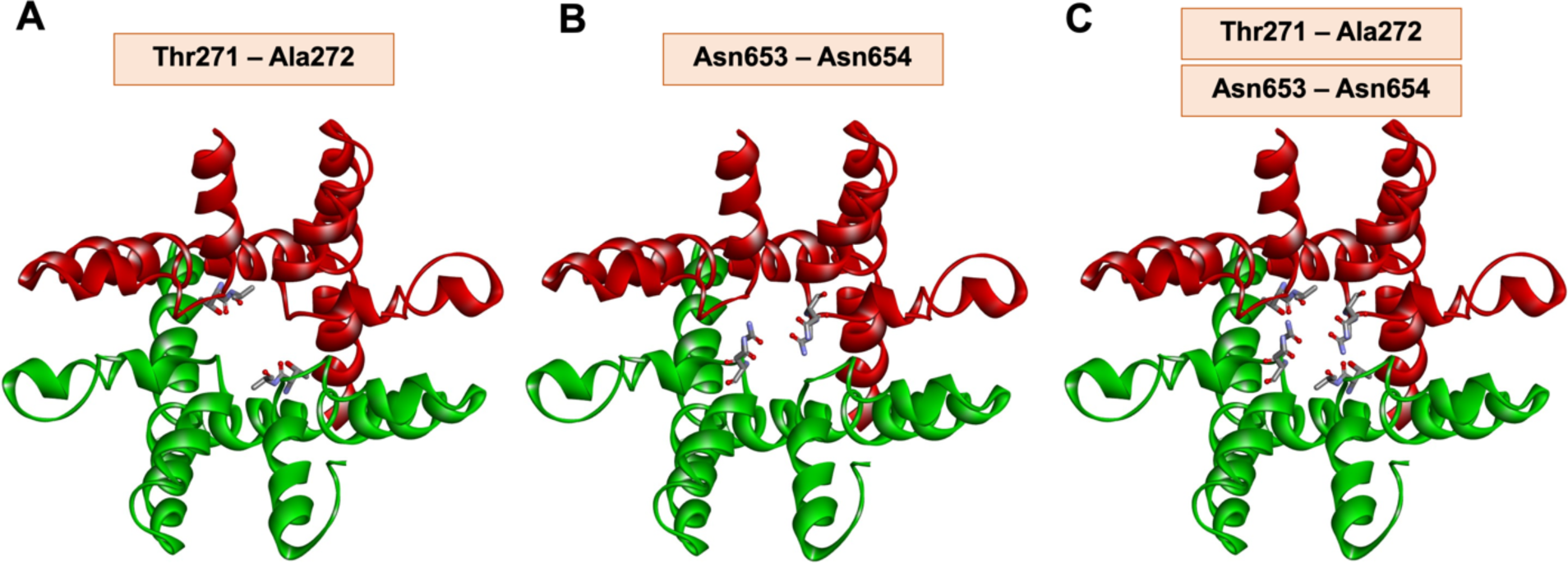
Top view of the previously suggested selectivity filters within the pore region. (A) Filter I central two residues; (B) filter II central two residues; (C) both filters shown together.

**Figure 4:**
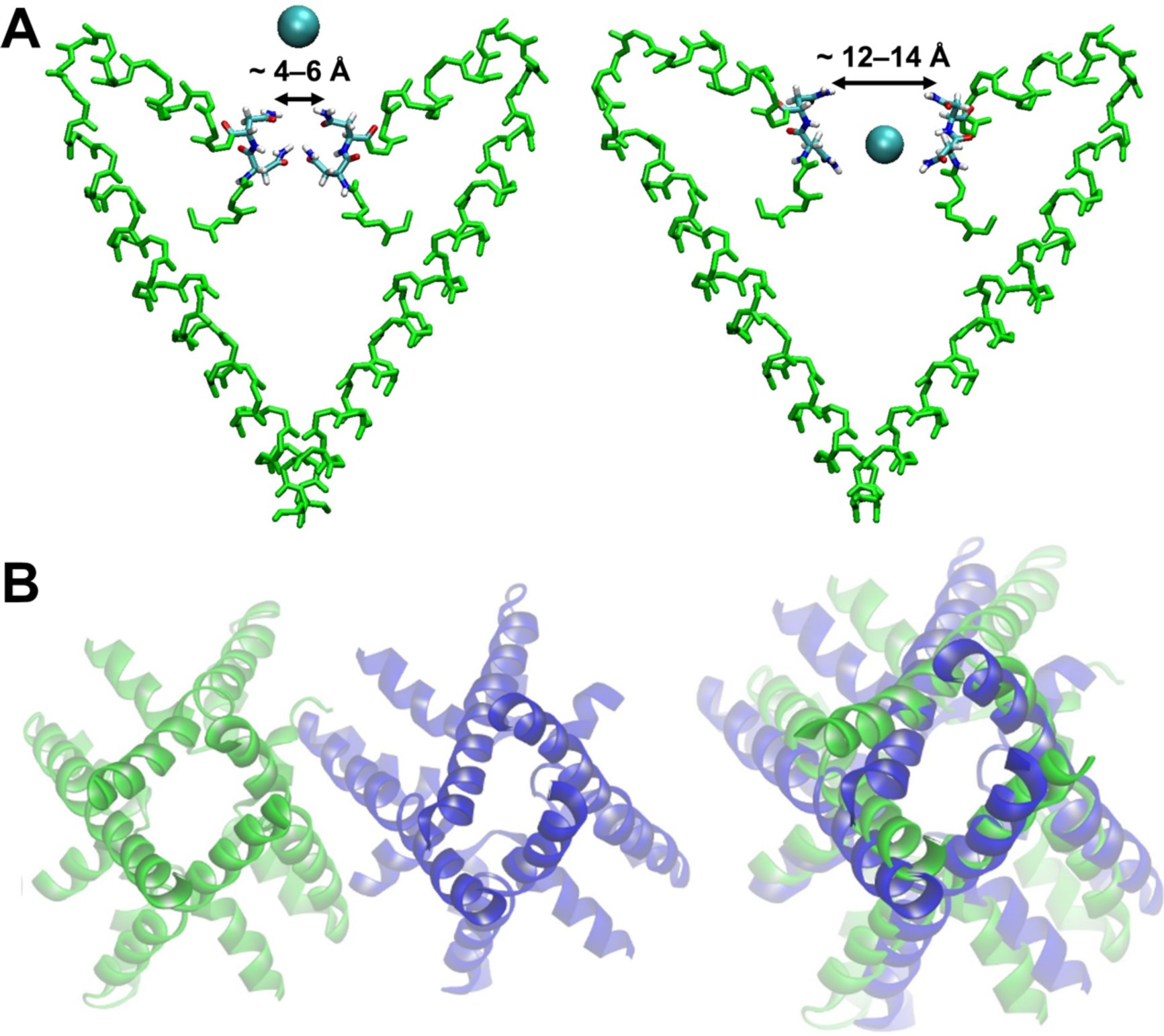
(A) Super-open state of the hTPC2 selectivity filter observed in our simulations together with the diameter of its filter entrance (right), compared to the selectivity filter geometry seen in the cryo-EM structure (PDB ID: 6NQ0; left). (B) Pseudo fourfold-symmetric open state of the hydrophobic gate at the cytoplasmic face of hTPC2 in simulations with PIP2 (green, left) and its transition towards a pseudo twofold-symmetric closed state without PIP2 (blue, centre). A superposition of both states highlights the constriction formed at the gate in the latter state (right).

Comparing our simulations of TPC2 with and without PIP2 further, we found that the apo form of TPC2 also adopted a closed state at the cytoplasmic hydrophobic gate, located at the bundle crossing of the channel (Fig. 4B). Whereas the hydrophobic gate in the PIP2-bound hTPC2 states became stabilised in an open, pseudo-C4 symmetric conformation, the apo hTPC2 states showed a transition towards a pseudo-C2 symmetric form, in which the minimum diameter of the hydrophobic gate (including side chains) was markedly reduced from 4 Å to 0 Å.

### Allosteric mechanism of PIP2-induced channel opening

To examine the coupling between ligand binding and pore dilation further, we first classed the TPC2 states into PIP2-bound super-open (ion-conducting), PIP2-bound closed (zero or only few ions traversing), and apo-closed states. We used principal component analysis (PCA) to identify the collective motions of the protein during the transition between the super-open and apo-closed states of the pore. The PCA eigenvectors were determined from a concatenated trajectory comprised of an equal number of frames in the PIP2-bound super-open and the apo-closed states. Subsequently, individual trajectories reflecting only super-open or apo-closed states were projected onto the eigenvectors obtained.

As can be seen in Fig. 5A, the projections onto eigenvectors 1 and 2 clearly separated the super-open (green) from the apo-closed (blue) states of the channel. The contribution of the first two principal components to the total variance was 49.7%, with the major distinction between open and closed states occurring on eigenmode 1. Analysing the correlated protein motion on eigenmode 1, we found that this mode represented opening and closing of the hydrophobic gate and the associated symmetry transition from pseudo-C4 to pseudo-C2 symmetry in the absence of PIP2. This motion of the pore-forming helical bundle was linked to a rotation of the VSDs, which contain the PIP2 binding sites, towards each other. More specifically, the PIP2-binding VSD of each subunit rotated towards the non-PIP2 binding VSD, again moving in a direction from pseudo-C4 to pseudo-C2 symmetry of the overall apo-protein structure (Fig. 5B). During this transition, an expansion of the space between S4, S4-S5 and S6, where the PIP2 binding site is located, was observed in the apo-state.

**Figure 5:**
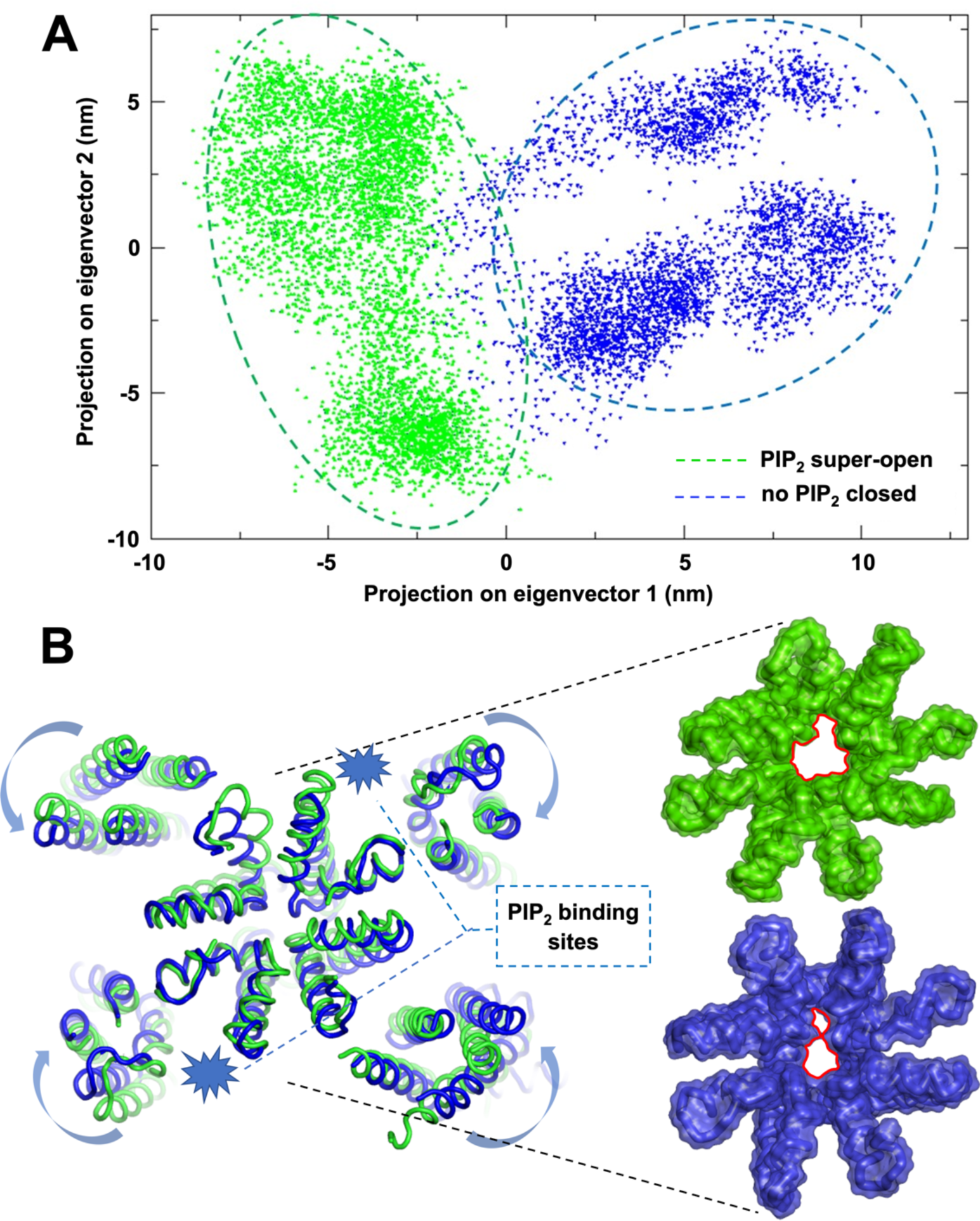
(A) Projection of the hTPC2 conformations found in simulations of the super-open, PIP2-bound state (green) and of the apo-closed state (blue), projected onto the first two eigenvectors of a PCA of the combined trajectories. (B) Concerted motion of the hTPC2 VSDs, linked to opening and closing of the SF and hydrophobic gate (left), and structures representing the super-open (green) and the apo-closed state (blue) projected on eigenmode 1 obtained from PCA. The reduction in gate diameter is highlighted in red.

### Ion conduction in TPC2

We next simulated hTPC2 under a range of ionic conditions and various different transmembrane voltages (V_m_, see Supplementary Material). Example permeation traces of individual Na^+^ ions along the pore axis, recorded in a simulation of a system with 0.6M NaCl at a V_m_ of -350 mV, are displayed in Fig. 6A. As can be seen, the major interacting site between the channel and the Na^+^ ions is observed in the central cavity of hTPC2. By contrast, and contrary to most cation channels, the luminal selectivity filter only showed transient interactions with the ions. The importance of the central cavity for ion conduction in hTPC2 has been recognised previously^[30]^; however, rather than forming a non-interacting aqueous volume occupied with a single ion, we found that a cluster of hydrophilic residues in the middle of the cavity (location marked in green in Fig. 6A) acted as a distinct binding site for Na^+^ ions. This cluster is composed of the conserved opposing asparagine residues N305 and N687 and the threonine residue T308 from each subunit (Fig. 6B). Additionally, we recorded interactions between the ions and the backbone oxygen atoms of V651 and V652 at the cavity-facing end of the SF.

**Figure 6:**
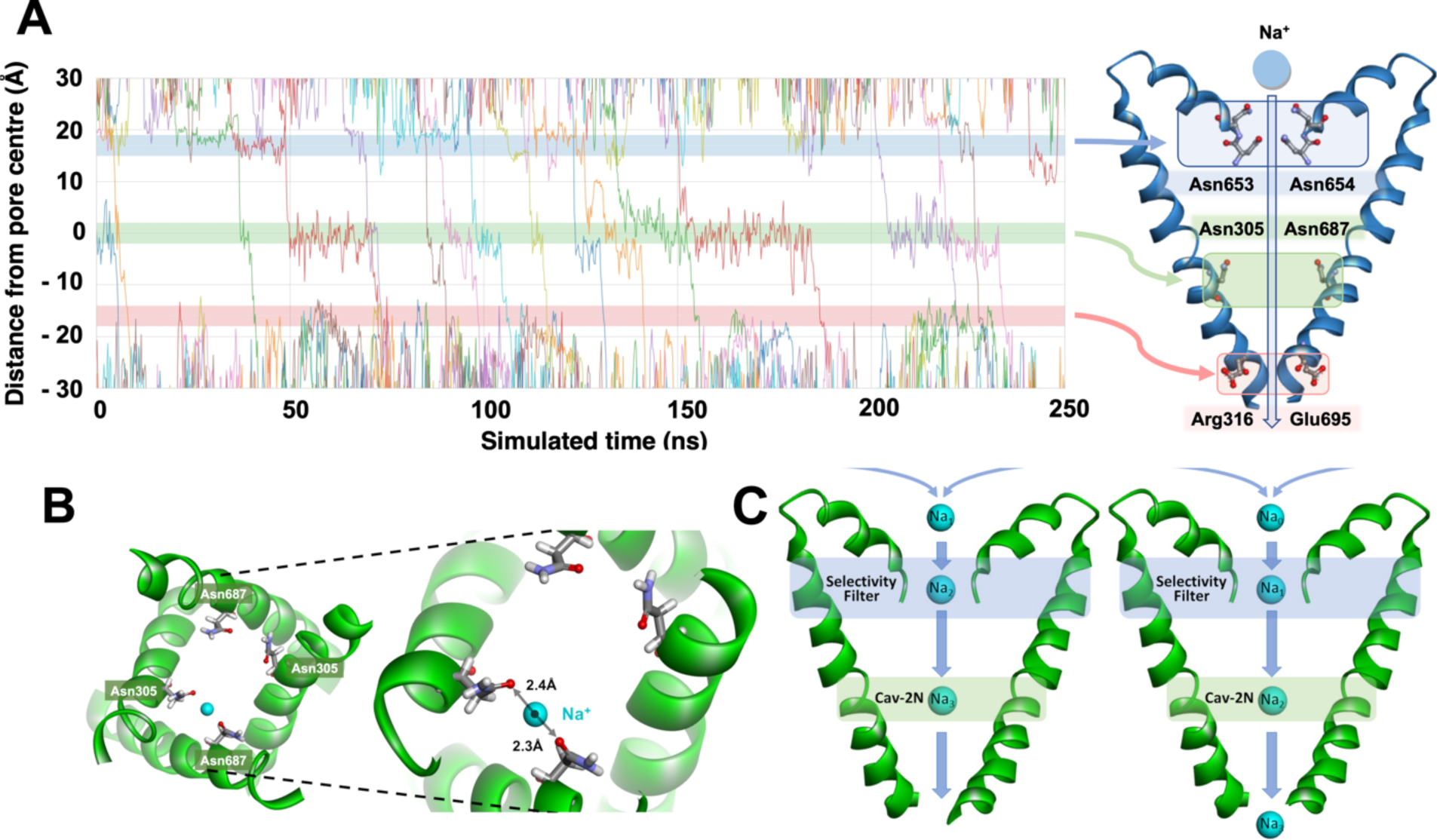
(A) Example traces of Na^+^ ions permeating hTPC2 show their intense interaction with the cavity hydrophilic cluster (green shade, residues N305 and N867 are highlighted), transient interactions with the SF (blue shade, residues N653 and N654 are highlighted), and fast permeation across the hydrophobic gate region (red shade; shown are the externally facing residues R316 and E695 near the gate for clarity). (B) Close-up view of residues N305 and N687 from two subunits, a Na^+^ ion binds between the asparagine side chains from one subunit but can change to other hydrophilic side chains in the cavity hydrophilic cluster. (C) Distant knock-on between a cavity-bound ion (Cav-2N) and an incoming ion transiently bound at the SF. The cavity serves as reservoir for Na^+^ ions.

In simulations of TPC2 in NaCl solutions, the Na^+^ ions interacted closely with the hydrophilic cluster over substantial time spans – it was occupied by Na^+^ during 75% of the simulated time on average, and the ions maintained an average minimum distance of 3.6 Å to the cluster-forming side chains in their bound state. Of note, the ions showed frequent transitions between the individual cluster-forming residues within the cavity, while remaining bound to the cluster as a whole. Conversely, the cations did not display any major interactions with the protein matrix during their traversal of the cytoplasmic hydrophobic gate region (Fig. 6A). A summary of all recorded ion permeation events in our simulations is shown in Table S1.

Additionally, Fig. 6A shows that a distant knock-on mechanism occurs between incoming ions passing through the SF and the ion stored over longer times within the central cavity. In this mechanism, the cavity-bound ion is pushed towards the cytoplasmic gate by the Coulombic interaction with the ion entering the cavity from the SF, which then replaces the previously bound ion (Fig. 6C). Due to the inherent permeation cooperativity of knock-on mechanisms, an increased level of efficiency is associated with this mechanism.

### The ion selectivity mechanism of TPC2

To study the ion selectivity mechanism of hTPC2, we next conducted five-fold repeated 500-ns simulations of mixed solutions composed of 0.3 mM NaCl and 0.3 mM CaCl_2_ at a V_m_ of -350 mV. In these simulations, the channels displayed a high selectivity for Na^+^ ions, with a permeability ratio of P_Na_:P_Ca_ = 9.4:1. Overall, we recorded 85 Na^+^ permeation events and 9 Ca^2+^ permeation events from the luminal entrance to the cytoplasmic exit of the channel during a total simulated time of 2.5 μs at a V_m_ of -350 mV. These findings are in very good agreement with the experimentally observed selectivity of P_Na_:P_Ca_ ≈ 10:1, reported from whole lysosomal recordings^[8]^.

The strict selectivity we found in our simulations raised the question how this selectivity is achieved in hTPC2, since both cation types did not exhibit strong interactions with the channel’s SF. Nonetheless, only few Ca^2+^ ions were seen to traverse the SF and subsequently enter the cavity. We therefore examined the favoured binding regions for each ion type in di-cationic NaCl and CaCl_2_ solution.

As shown in Fig. 7A, the ion density maxima found for Na^+^ ions near the protein matrix are located on the pore axis, within the central cavity, and throughout the VSD. In contrast, on the luminal side of the channel, the main Ca^2+^ binding sites occur off the pore axis near acidic clusters consisting of residues D276, D660, and E260. The Ca^2+^ density maxima are not observed near the channel entrance on the pore axis, the SF, or the central cavity, such that the local Ca^2+^ concentration is depleted compared to the local Na^+^ concentration at the luminal entrance to hTPC2. All Ca^2+^ ions that did enter the cavity first bound to the same hydrophilic cluster region within the cavity as the Na^+^ ions at N305, N687 and T308, and subsequently fully permeated the channel, driven by V_m_. Fig. 7B shows the permeation traces of all permeating cations in five-fold repeated simulations of hTPC2 under di-cationic conditions at a V_m_ of -350 mV.

**Figure 7:**
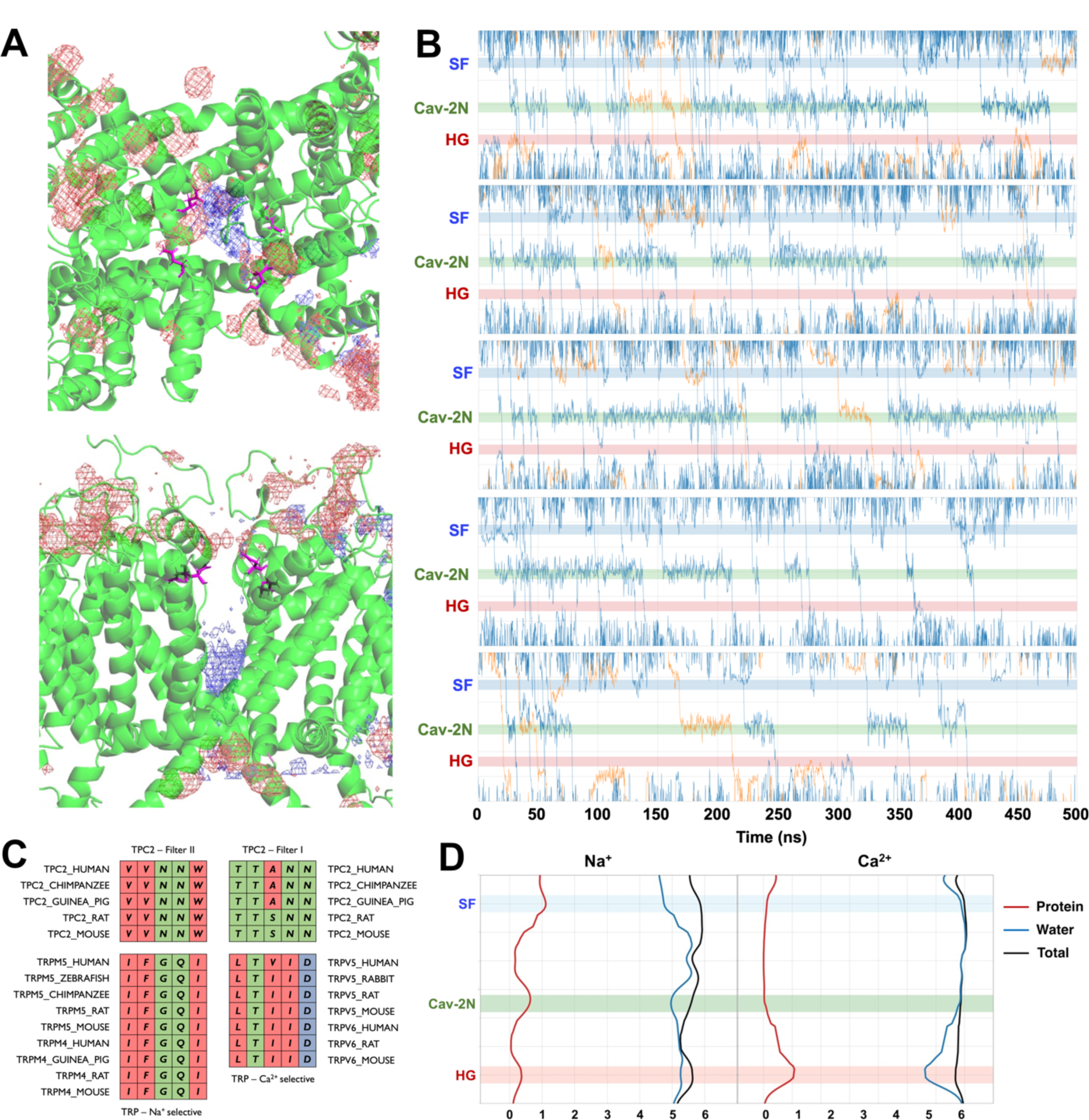
(A) Density of Ca^2+^ ions (red mesh) and Na^+^ ions (blue mesh) in simulations around hTPC2 performed in equimolar di-cationic solution. Top: View from the lumen; bottom: side view. (B) Permeation traces of Na^+^ (blue) and Ca^2+^ ions (orange) across hTPC2 in equimolar di-cationic solution at V_m_ = -350 mV demonstrate the channel’s selectivity for Na^+^. From the luminal to the cytoplasmic side, the SF, cavity hydrophilic cluster (Cav-2N), and hydrophobic gate (HG) are shaded blue, green, and red, respectively. (C) Comparison of the hTPC filter I and II amino acid sequences with the SF sequences of related monovalent-selective and Ca^2+^ selective TRP channels. (D) Interactions of Na^+^ and Ca^2+^ ions with water oxygen atoms (blue) and protein oxygen atoms (red) within their first solvation shell. Na^+^ ions show closer interactions with the protein, especially at the SF and in the central cavity.

In addition to the lowered local concentration at the SF, Ca^2+^ ions were typically not able to pass through the SF and the transition region between the SF and the cavity. The sequence of both hTPC2 filters I and II contains a pair of neutral asparagine residues instead of a negatively charged aspartate residue, where aspartates are often related to tight Ca^2+^ interactions and are present, for example, in the SFs of the Ca^2+^ selective channels TRPV5 and TRPV6 (Fig. 7C). By contrast, the sequence of filter II resembles the SF sequences of TRPM4 and TRPM5 channels, which are both highly selective for monovalent cations *vs.* Ca^2+[47]^. In our simulations, we observed that permeating cations interact more closely with filter II than with filter I. For instance, close contacts between the ions and filter residues with distances lower than 3 Å are observed 2-3 times more frequently with filter II. As shown by the solvation profile of both ion types (Fig. 7D), the SF matrix replaces one water molecule in the hydration shell of Na^+^ through transient interactions, especially with the asparagine side chains of filter II (N653 and N654). In contrast, these interactions are absent in the case of Ca^2+^ ions, which permeate the SF in a fully hydrated state.

Furthermore, the SF-cavity transition zone is formed by the hydrophobic residue pair V651 and V652 in filter II, whereas a pair of polar threonine residues, T270 and T271, line this region in filter I. Our previous simulations on the TRPM5 channel suggested that traversing such a hydrophobic tunnel region incurs a lower energy penalty for monovalent cations than for divalent cations due to the difference in Born energy _[41,48,49]_. Since the ions permeate mainly along filter II, this hydrophobic region is thus likely to contribute further to the selectivity of hTPC2 for the monovalent Na^+^ ions.

Finally, a polar cluster within the central cavity (consisting of N305, N687, and T308) shows strong interactions with Na^+^ ions, leading to a particularly long residence time of Na^+^ ions in the cavity. The cavity thereby acts as a reservoir for ions, which supports an efficient mechanism where a distant knock-on leads to the collective motion of an incoming ion with the cavity ion, whereby the cavity ion traverses the hydrophobic gate to the cytoplasm and is replaced by the incoming ion.

Our simulations therefore suggest a multi-layered mechanism of Na^+^ selectivity in hTPC2, combining the luminal face of the channel, the SF, and the central cavity. At the luminal channel entrance, the Ca^2+^ concentration is locally depleted due to strong binding of the divalent Ca^2+^ ions to the negatively charged Asp276, Asp660, and Glu260 side chains distal to the SF entrance. Within the SF itself, no charged residues are present which could bind Ca^2+^ ions, whereas the filter II residues N653 and N654 show transient interactions only with Na^+^ ions. At the entrance to the central channel cavity, a region of raised hydrophobicity in filter II, at V651 and V652, adds a further layer of selectivity for monovalent cations. Moreover, the long residence time of Na^+^ ions at a hydrophilic cluster within the cavity facilitates an efficient cooperative permeation mechanism for Na^+^ ions by distant knock-on in combination with an incoming ion.

### Structural and functional importance of the disulfide bond between Cys623 residues

Within the second domain of the hTPC2 homodimers, the S5 helices and the pore-constructing helices P1-P2 are connected by two lumen-facing external loops (Fig. 8). These two loops consist of 37-38 residues and are interlinked via a disulfide bond between the C623 residues of each subunit, which are specific to hTPC2. Similar cysteine residues are absent from the corresponding regions in mTPC1 and *at*TPC1. We were therefore interested if the inter-subunit disulfide bond played an additional functional role. We simulated a C623A mutant and compared the results to simulations of the wild-type containing the disulfide bond.

**Figure 8:**
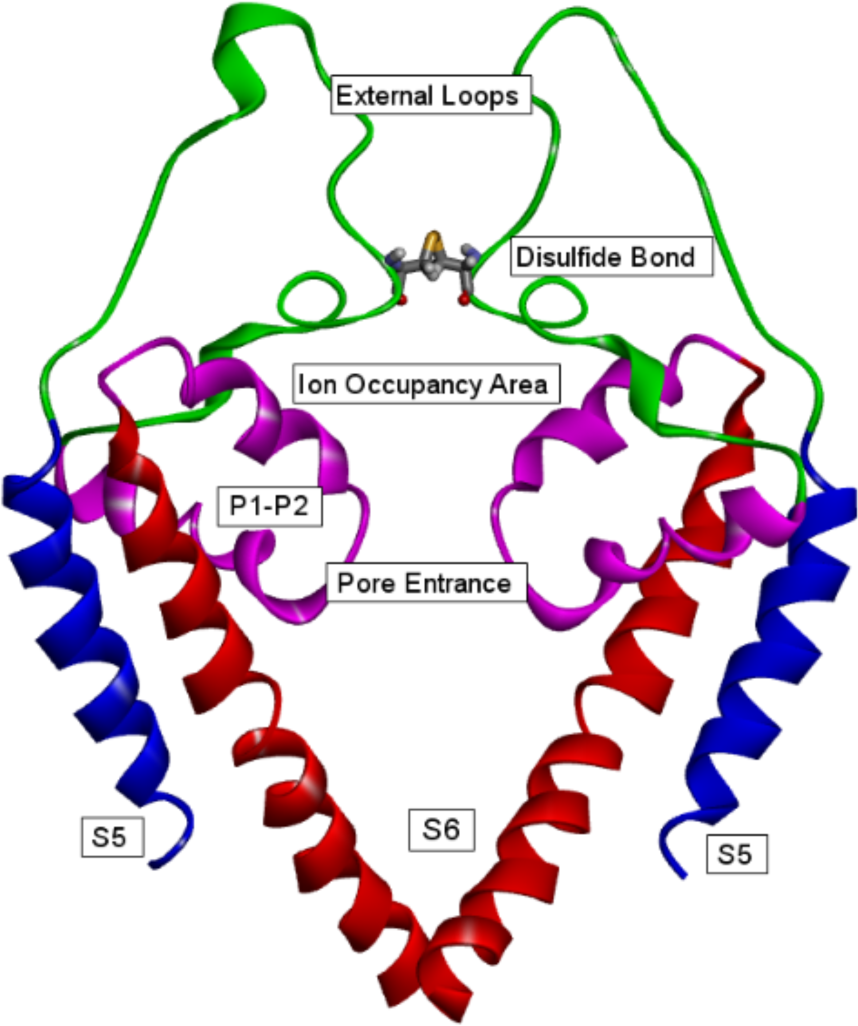
Pore and external loop region of hTPC2. The disulfide bond between C623 from each subunit maintains an accessible pore entrance and rigidifies the luminal face of hTPC2.

In 5-fold repeated 500-ns long simulations of C623A hTPC2, we did not observe any permeation events for Na^+^ – the most highly permeable ion in hTPC2. Whereas in the wild-type, the disulfide bond acted as an upper roof for the ion recruitment area (see Fig. 9), preventing the collapse of the long external loops into either this area or further interactions near the pore-forming domains, the loops folded back inwards and closed the pore from the luminal side in the mutant. Close to the C623 residue, charged and polar residues such as E630, Q628, E627, S625, S620 and N618 were observed to become more flexible in the mutant in the absence of the C653 disulfide bridge. These residues strongly interact with each other in the C653A mutant, forming a physical barrier to ion permeation at the entrance of the pore. According to our simulations, the C623 disulfide bond is therefore important in keeping the luminal channel entrance in a conformation accessible to ions.

## Conclusion

TPCs possess two different amino acid sequences forming their SFs. As shown by Guo et al.^[28]^, mutations in filter II exert a greater effect on ion selectivity of TPC2 than mutations in filter I. Our simulations reproduce the Na^+^ selectivity of PIP2 and demonstrate that especially the asparagine residues in filter II interact strongly, albeit transiently, with Na^+^ ions, forming the central element of a three-level selectivity mechanism, which includes the luminal entrance, SF, and the central channel cavity. The role of the asparagine residues in the SF is mirrored by a cluster of polar residues including two asparagine residues in the cavity. This binding site within the cavity is almost continually occupied by a Na^+^ ion, which enables a distant knock-on mechanism to occur between any incoming ion at the SF and the cavity-bound ion, enhancing the conduction efficiency for Na^+^ ions.

Simulations with and without PIP2 show that PIP2 binding not only allosterically opens the hydrophobic cytoplasmic gate of the channel, but also leads to a further opening of the SF, resulting in a high-conductance conformation of TPC2 with a wider SF than seen in the cryo-EM structure^[16]^. Our simulations indicate that this may be a functionally relevant state, as without this additional widening ion conduction occurs only very rarely. Finally, the geometry of the loops at the luminal entrance of TPC2 plays an important role in conduction, as a disulfide bridge appears to be necessary to allow ions to first access the SF.

## Acknowledgements

We thank Callum Ives for assistance with the simulations analysis and useful discussions and Samantha Pitt for drawing our attention to the TPC channel family, helpful discussions, and careful reading of the manuscript. Alp Tegin Şahin was supported by an EASTBio PhD studentship from the Biotechnology and Biological Sciences Research Council (BBSRC) BB/T00875X/1.

