## Supplementary Material for "Characterisation of ligand gating, ion conduction and the ion selectivity mechanism in the endo-lysosomal ion channel hTPC2"

*Table S1: Overview of the simulated systems, including applied voltage (mV), number of repeats, total simulation time and number of ion permeation events.*

| System | Applied Voltage (mV) | Number of Repeats | Total Simulation Time (μs) | Number of Permeation Events |
|---|---|---|---|---|
| POPC / 0.6M NaCl | + 700 | 5 | 1.75 | 152 $Na^+$ |
| | - 700 | 5 | 1.75 | 76 $Na^+$ |
| | - 350 | 5 | 1.75 | 22 $Na^+$ |
| | - 200 | 5 | 1.75 | 0 |
| POPC / 0.6M $CaCl_2$ | - 700 | 5 | 1.75 | 4 $Ca^{2+}$ |
| WO/PIP2 | -700 | 3 | 1 | 5 $Na^+$ |
| WO/PIP2 | -200 | 3 | 1 | 0 $Na^+$ |
| LysM / 0.6 NaCl | - 700 | 5 | 1.75 | 49 $Na^+$ |
| LysM / 0.6 $CaCl_2$ | - 700 | 5 | 1.75 | 0 |
| POPC / 0.3M NaCl&$CaCl_2$ | - 700 | 5 | 5 | 122 $Na^+$ / 66 $Ca^{2+}$ |
| POPC / 0.3M NaCl&$CaCl_2$ | - 350 | 5 | 2.5 | 70 $Na^+$ / 9 $Ca^{2+}$ |
| C623A Mutant | -700 | 5 | 2.5 | 1 $Na^+$ |
| **Total** | | **56** | **24.25 μs** | **497 $Na^+$ / 79 $Ca^{2+}$** |

*Table S2: Lipid composition of mammalian lysosomal membranes.*

| * LYSm (mammalian lysosomal membranes) | | | | |
|---|---|---|---|---|
| Lipid Name | Lipid Head/Tail | #Lipids in Leaflets | APL( ) | Charge(*e*) |
| | | Outer and Inner | Inner and Inner | |
| PLPC | PC(16:0/18:2(9Z,12Z)) | 11 | 62.43 | 0 |
| SAPC | PC(18:0/20:4(5Z,8Z,11Z,14Z)) | 18 | 64.47 | 0 |
| SDPC | PC(18:0/22:6(4Z,7Z,10Z,13Z,16Z,19Z)) | 6 | 63.3 | 0 |
| PLPE | PE(16:0/18:2(9Z,12Z)) | 7 | 62.01 | 0 |
| SAPE | PE(18:0/20:4(5Z,8Z,11Z,14Z)) | 18 | 54.26 | 0 |
| SAPI | PI(18:0/20:4(5Z,8Z,11Z,14Z)) | 6 | 63.63 | -1 |
| SDPI | PI(18:0/22:6(4Z,7Z,10Z,13Z,16Z,19Z)) | 2 | 66 | -1 |
| SAPS | PS(18:0/20:4(5Z,8Z,11Z,14Z)) | 2 | 65.98 | -1 |
| SDPS | PS(18:0/22:6(4Z,7Z,10Z,13Z,16Z,19Z)) | 1 | 61.05 | -1 |
| PSM | SM(d18:1/16:0) | 3 | 59.53 | 0 |
| LSM | SM(d18:1/24:0) | 3 | 56.65 | 0 |
| CHOL | Cholesterol | 18 | 30.68 | 0 |
| BMP | BMP(18:1(9Z)/18:1(9Z)) | 7 | 76.13 | -1 |
